# *Pneumocystis murina* Promotes Inflammasome Formation and NETosis during *Pneumocystis* Pneumonia

**DOI:** 10.1101/2024.02.16.580773

**Authors:** Steven G. Sayson, Alan Ashbaugh, Aleksey Porollo, Melanie T. Cushion

## Abstract

*Pneumocystis jirovecii* pneumonia (PjP) poses a serious risk to individuals with compromised immune systems, such as individuals with HIV/AIDS or undergoing immunosuppressive therapies for cancer or solid organ transplants. Severe PjP triggers excessive lung inflammation, resulting in lung function decline and consequential alveolar damage, potentially culminating in acute respiratory distress syndrome. Non-HIV patients face a 30-60%mortality rate, emphasizing the need for a deeper understanding of inflammatory responses in PjP.

Prior research emphasized macrophages in *Pneumocystis* infections, neglecting neutrophils’ role in tissue damage. Consequently, the overemphasis on macrophages led to an incomplete understanding of the role of neutrophils and inflammatory responses. In the current investigation, our RNAseq studies on a murine surrogate model of PjP revealed heightened activation of the NLRP3 inflammasome and NETosis cell death pathways in their lungs. Immunofluorescence staining confirmed Neutrophil Extracellular Trap (NET) presence in the lungs of the *P. murina*-infected mice, validating our findings. Moreover, isolated neutrophils exhibited NETosis when directly stimulated with *P. murina*. While isolated NETs did not compromise *P. murina* viability, our data highlight the potential role of neutrophils in promoting inflammation during *P. murina* pneumonia through NLRP3 inflammasome assembly and NETosis. These pathways, essential for inflammation and pathogen elimination, bear the risk of uncontrolled activation leading to excessive tissue damage and persistent inflammation.

This pioneering study is the first to identify the formation of NETs and inflammasomes during *Pneumocystis* infection, paving the way for comprehensive investigations into treatments aimed at mitigating lung damage and augmenting survival rates for individuals with *PjP*.

**IMPORTANCE:** *Pneumocystis jirovecii* pneumonia (PjP) affects individuals with weakened immunity, such as HIV/AIDS, cancer, and organ transplant patients. Severe PjP triggers lung inflammation, impairing function and potentially causing acute respiratory distress syndrome. Non-HIV individuals face a 30-60% mortality rate, underscoring the need for deeper insight into PjP’s inflammatory responses.

Past research focused on macrophages in managing *Pneumocystis* infection and its inflammation, while the role of neutrophils was generally overlooked. In contrast, our findings in *P*. *murina*-infected mouse lungs showed neutrophil involvement during inflammation and increased expression of NLRP3 inflammasome and NETosis pathways. Detection of neutrophil extracellular traps further indicated their involvement in the inflammatory process. Although beneficial in combating infection, unregulated neutrophil activation poses a potential threat to lung tissues.

Understanding the behavior of neutrophils in *Pneumocystis* infections is crucial for controlling detrimental reactions and formulating treatments to reduce lung damage, ultimately improving the survival rates of individuals with PjP.

## INTRODUCTION

*Pneumocystis jirovecii* pneumonia (PjP) is a leading cause of mortality in hospitalized individuals with HIV/AIDS. However, the incidence of PjP has increased in cancer patients and individuals who have received organ transplants requiring immune-suppressing treatments. Among all hospitalizations for PjP, malignancy stands as the most prevalent predisposing factor, accounting for 46.0 to 55.7% of cases, followed by HIV at 17.8% (1–4). Within immunocompetent hosts, signaling cascades initiated by CD4+ T cells and B cells efficiently trigger *P. jirovecii* clearance with minimal inflammation (5). Conversely, immunosuppressed hosts experience an influx of various immune cells into their lungs, including T lymphocytes, polymorphonuclear neutrophils (PMNs), and other leukocytes (6–9). This can lead to a profound inflammatory response which leads to considerable morbidity and mortality. Severe PjP is characterized by a neutrophilic inflammatory response presenting with decreased pulmonary function, alveolar damage, and respiratory failure (10, 11). Furthermore, Bronchial alveolar lavage (BAL) neutrophilia has been shown to be a predictor of poor prognosis and increased mortality in PjP (12).

Previous research has primarily centered on macrophage polarization and its role in clearance of *Pneumocystis* from infected hosts. Both classically activated M1 and alternatively activated M2 macrophages are involved in *Pneumocystis* clearance. Immunocompetent mice show preferential activation of the M2 phenotype during *P. murina* exposure, while immunosuppressed hosts show enhanced M1 polarization (13). M1-polarized macrophages are effective fungicidal cells and produce a substantial cytokine and chemokine response (14). These secretions not only eliminate infectious organisms but also signal for further recruitment of immune cells.

Pattern recognition receptors (PRRs), such as C-type lectin receptors Dectin-1/2 and toll-like receptors (TLRs), expressed by innate immune cells bind to pathogen-associated molecular patterns (PAMPs) and damage-associated molecular patterns (DAMPs) (15). Dectin-1/2 and TLR2 have been identified as important receptors in antigen recognition to *P. murina* in mice, leading immune responses such as pro-inflammatory cytokine release and fungal clearance (16–18).

Neutrophils play a crucial role in host defenses against pathogenic organisms. During *Candida albicans* and *Aspergillus fumigatus* infections in mice, neutrophils aid in controlling fungal growth by phagocytosis and ROS generation for effective pathogen killing (19, 20). Beyond phagocytosis and ROS generation, neutrophils employ additional antimicrobial activity, including degranulation, cytokine production, and NETosis. NETosis results in the expelling of DNA to form <u>n</u>eutrophil <u>e</u>xtracellular traps (NETs) that ensnare and kill pathogens (21, 22). These NETs are decorated with various proteins, such as histones, neutrophil elastase (NE), calprotectin, myeloperoxidase (MPO), and other antimicrobial proteins (23, 24).

A previous study by Swain et al. found that neutrophils and reactive oxygen species (ROS) do not contribute to pulmonary tissue damage, nor play a major role in clearance of *Pneumocystis* (25). Consequently, despite the significant influx of neutrophils into the lungs during *Pneumocystis* infection, their role in inflammation has been both overlooked and understudied, resulting in a significant knowledge gap.

This study revealed an upregulation in the expression of genes associated with the NLRP3 inflammasome and NETosis in the lungs of mice infected with *P. murina*. These processes are essential for the elimination of pathogens from the host system. However, overactivation of these pathways leads to heightened inflammation and tissue damage, emphasizing the need for a thorough understanding of their regulation in the context of *Pneumocystis* infection. This investigation intended to elucidate the roles of neutrophil populations during *PjP* to establish the groundwork for therapeutic strategies aimed at mitigating excessive inflammatory responses and alveolar damage.

## RESULTS

### Signaling pathways involved with inflammatory processes increase during *P. murina* infection

To gain a better understanding of the changes in host immune response during the development of *P. murina* infection in mice, lung samples were analyzed from both uninfected and *P. murina*-infected immunosuppressed mice at 5- and 7-week intervals after initial exposure to *P. murina*. Mice with a 7-week infection exhibited a higher fungal burden than those with 5-week infections (Figure 1A). Differential gene expression analysis revealed that gene expression patterns across different samples were comparable between biological groups (Figure 1B). Control samples from both the 5-week (MC) and 7-week (HC) groups clustered at the bottom, while infected samples from the 5-week (MI) group clustered in the middle, and those from the 7-week (HI) group clustered at the top of the heatmap. A gene set enrichment analysis indicated a significant increase in signaling pathways associated with inflammatory processes as *P. murina* infection progressed in mice (Figure 1C; Table 1).

**Figure 1.**
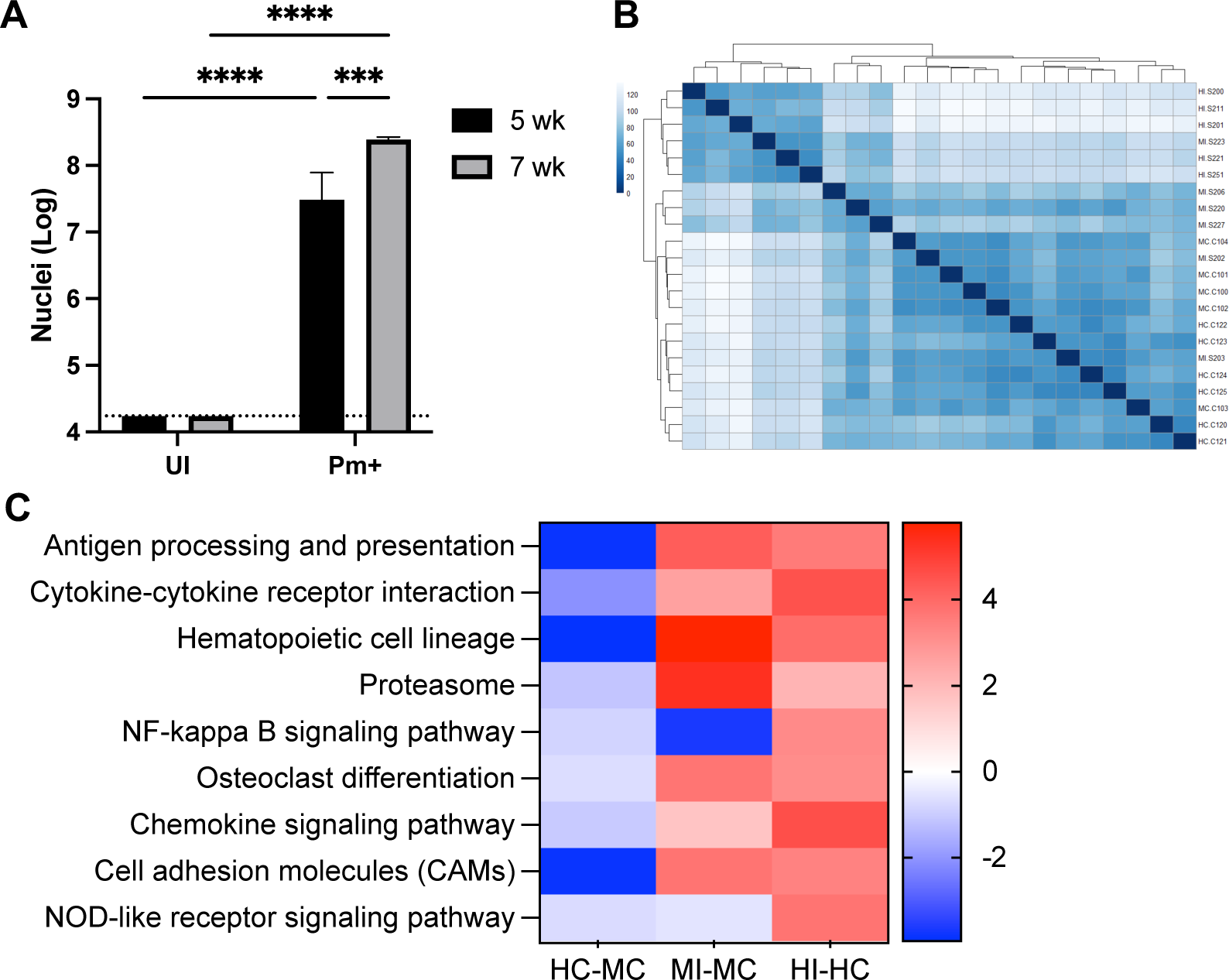
Signaling pathways associated with inflammatory processes are up-regulated as *P. murina* infection progressed in mice. Immunosuppressed mice were exposed to uninfected or previously *P. murina*-infected mice. (A) 5- or 7-weeks after exposure, the fungal burden in the lungs were enumerated. Dashed line, limit of microscopic enumeration. *, *p<0.05.* (B) Heatmap of sample-to-sample distances. 5-week (MC) and 7-week (HC) control, uninfected samples cluster near the bottom. 5-week infected (MI) cluster in the middle, while 7-week infected (HI) cluster at the top of the heatmap. (C) Heatmap comparison analysis of Gene Set Enrichment Analysis on KEGG Pathways relating to signaling. Data is represented as logFC.

**Table 1.** Functional gene set enrichment analyses. Enriched pathways based on differential expression.

| Pathway | KEGG ID | NumGenes<sup>a</sup> | p-value | FDR |
| --- | --- | --- | --- | --- |
| Antigen processing and presentation | mmu04612 | 77/91 | 1.37E-17 | 3.25E-16 |
| Cytokine-cytokine receptor interaction | mmu04060 | 222/270 | 3.21E-23 | 3.06E-21 |
| Hematopoietic cell lineage | mmu04640 | 83/95 | 2.26E-12 | 2.81E-11 |
| Proteasome | mmu03050 | 45/45 | 9.61E-11 | 9.82E-10 |
| NF-kappa B signaling pathway | mmu04064 | 89/103 | 2.38E-15 | 4.53E-14 |
| Osteoclast differentiation | mmu04380 | 116/130 | 1.45E-10 | 1.38E-09 |
| Chemokine signaling pathway | mmu04062 | 162/192 | 2.30E-08 | 1.68E-07 |
| Cell adhesion molecules (CAMs) | mmu04514 | 140/169 | 4.80E-15 | 8.57E-14 |
| NOD-like receptor signaling pathway | mmu04621 | 152/169 | 1.54E-08 | 1.22E-07 |

<sup>a</sup> NumGenes, number genes mapped to the total number of genes in the set

### NOD-like signaling and NETosis pathways are increased during *P. murina* **infection.**

Many of the inflammatory signaling pathways during *Pneumocystis* infections have been extensively studied, such as antigen processing and presentation, cytokine-cytokine receptor interaction, hematopoietic cell lineage, NF-kappa B signaling, chemokine signaling pathway, and cell adhesion molecules (26–30). For that reason, we chose to focus on the understudied NOD-like receptor signaling pathway. In the lungs of mice with both 5- and 7-week infections, there was an observed increase in the expression of key genes associated with the NOD-like receptor signaling pathway that were not present in non-infected mice. These genes included NOD-like protein receptor 3 (*Nlrp3*), Apoptosis-associated Speck-like protein containing a CARD (*Asc/Pycard*), and caspase 1 (*Casp1*) (Figure 2). These transcripts encode proteins that form the NLRP3 inflammasome complex, wherein NLRP3 binds to the scaffold protein ASC, which in turn binds to CASP1. The proteolytic enzyme CASP1 component of the NLRP3 inflammasome complex cleaves the pro-inflammatory cytokine interleukin 1 beta (IL-1β) into its mature form. Additionally, CASP1 also cleaves gasdermin D (GSDMD), which inserts into the plasma membrane to facilitate the release of IL-1β to further propagate a proinflammatory response and pyroptotic cell death (31).

**Figure 2.**
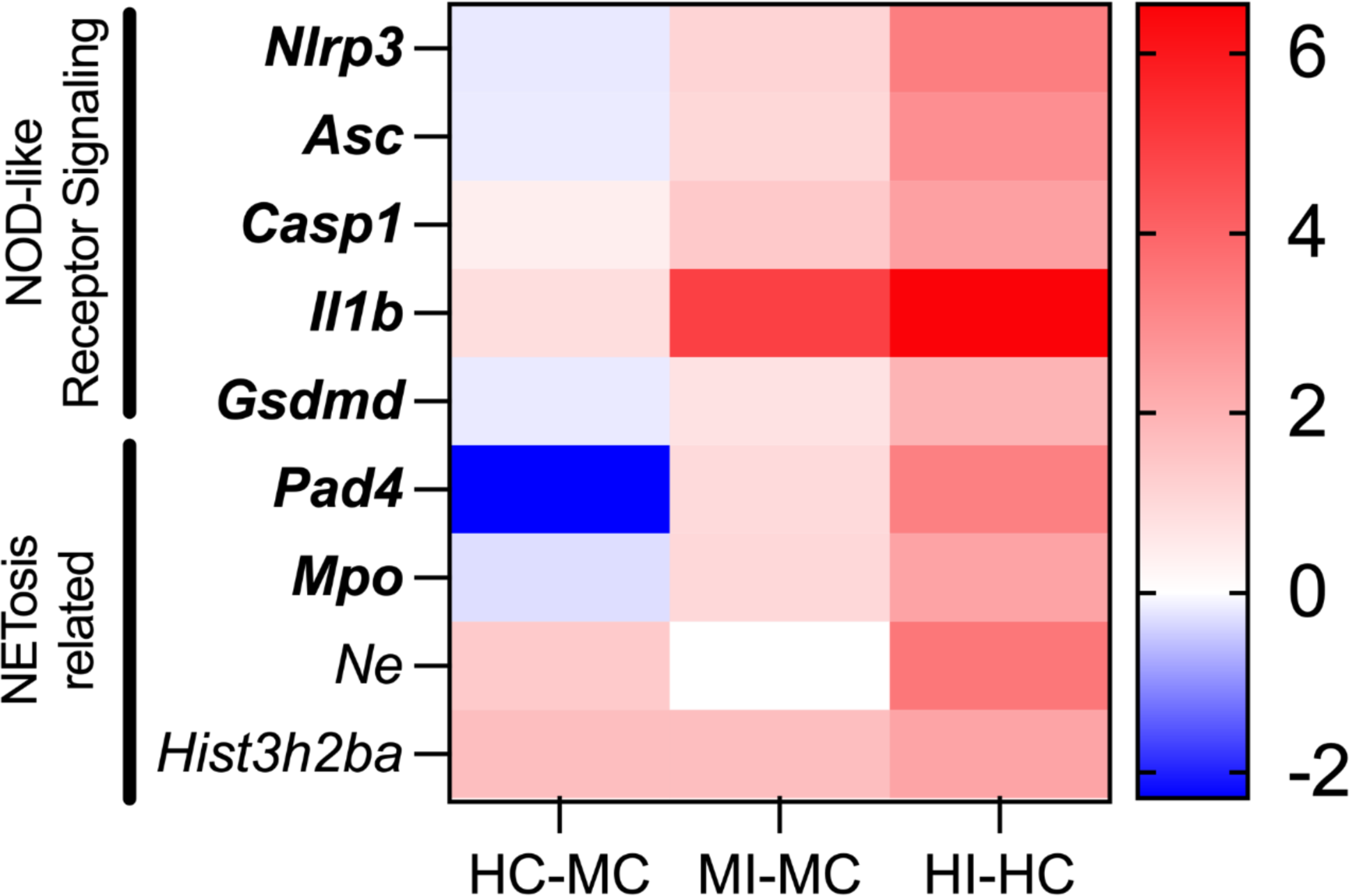
NOD-like receptor signaling and NETosis related transcripts are up-regulated as *P. murina* infection progressed in mice. Data is represented as log2-fold change between the displayed groups. 5-week uninfected, MC. 7-week uninfected, HC. 5-week infected, MI. 7-week infected, HI. Data is represented as logFC. Bold, FDR adjusted p<0.05.

The mRNA expression of both *Il-1β* and *Gsdmd* increased in the lungs of mice with 5-week and 7-week infections (Figure 2). While proinflammatory cytokine IL-1 is a crucial mediator for host resistance and immune cell recruitment during *P. murina* infection (32), inflammasome assembly details have yet been described. Additionally, increased peptidylarginine deiminase 4 (*Pad4*) expression was detected in *P. murina*-infected mice. PAD4 plays a role in chromatin decondensation and expulsion of NETs in a regulated cell death pathway called NETosis (33). Furthermore, PAD4 stimulates NLRP3-inflammasome formation, leading to prolonged inflammation through a positive feedback loop by driving the production of pro-inflammatory cytokines and chemokines (34). This is supported by similar findings that bronchial cells exposed to NETs demonstrated increased secretion of pro-inflammatory cytokine IL-1β (35).

### NET Formation in *P. murina*-Infected Lung Tissue

Given the increased expression of genes related to NOD-like receptor signaling and the NETosis pathway through differential expression analysis, we sought to confirm the presence of NETs during *Pneumocystis murina* pneumonia (PmP). Increased expression of NE and MPO, which decorate the extracellular DNA in NETs, are significantly increased in the lungs of *P. murina*-infected mice (Figure 3A). Immunofluorescence analysis revealed the expression of citrullinated histone H3 (CitH3) and NE, suggesting the presence of NETs within *P. murina*-infected lung tissue (Figure 3B).

**Figure 3.**
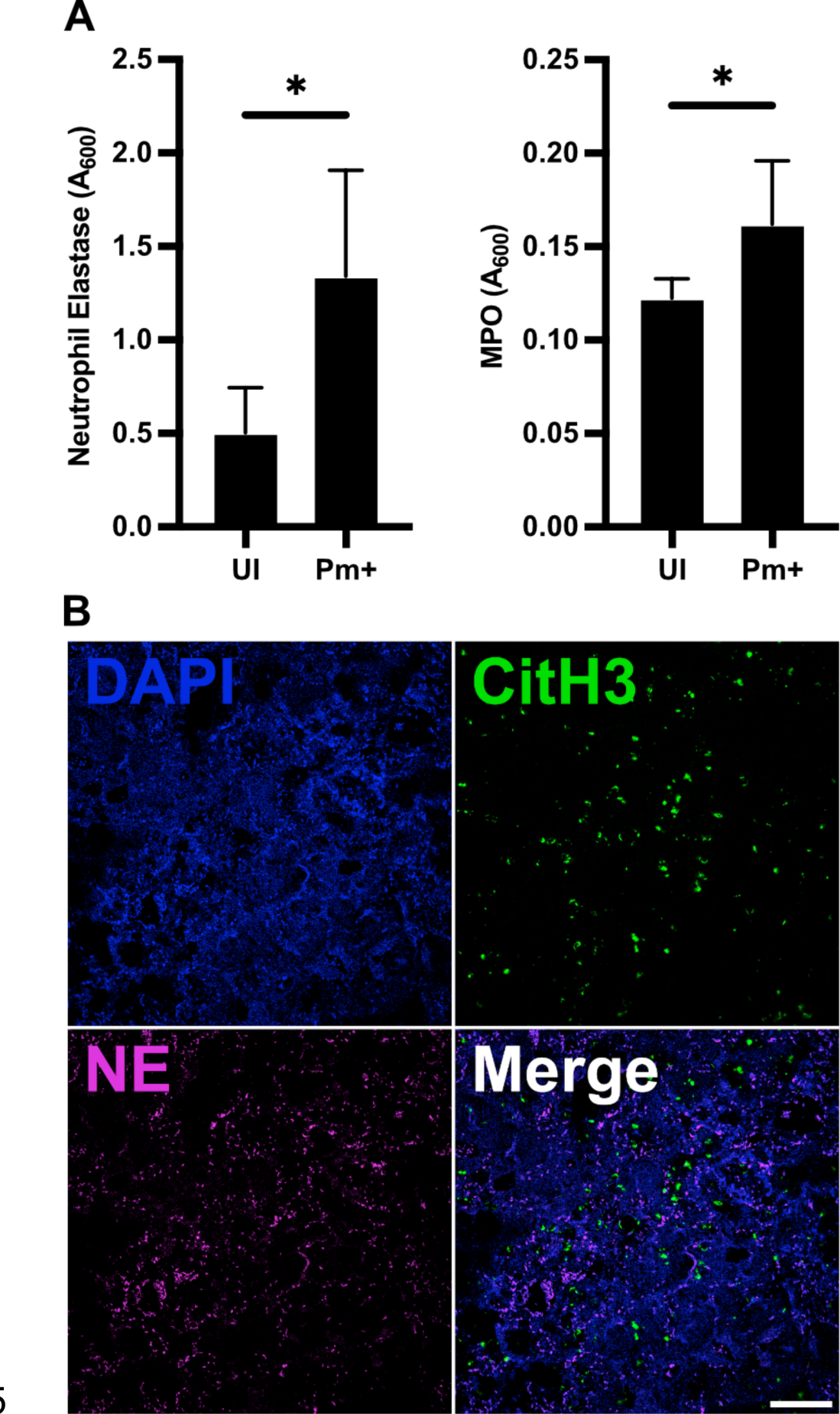
NETosis-related proteins are present in the lungs of *P. murina*-infected mice. (A) ELISA reveals increased expression of NET components, neutrophil elastase and myeloperoxidase (MPO) in the lung tissue from *P. murina*-infected mice. *t-test, *p*<0.05. (B) Immunofluorescence of the lungs of *P. murina*-infected mice. The NETosis markers, neutrophil elastase (green) and citrullinated histone H3 (yellow), are present throughout the lung. DAPI, blue. Images taken on a Nikon A1 confocal microscope.

### *P. murina* stimulates NETosis in neutrophils *in vitro*

Considering that *Pneumocystis* infections result in lung neutrophilia and NETs were observed within the lungs of *P. murina*-infected mice, we investigated whether *P. murina* could directly induce neutrophils to undergo NETosis and produce NETs. In experiments using bone marrow-derived neutrophils, we found that *P. murina* stimulated the release of NETs in neutrophils, as evidenced by the extracellular DNA released into the supernatant in a dose-dependent manner (Figure 4A). Additionally, since NETs carry various proteins, we conducted a sandwich ELISA to detect MPO-DNA complexes within the NETs, which showed that *P. murina* stimulated the release of DNA-MPO complexes from neutrophils (Figure 4B).

**Figure 4.**
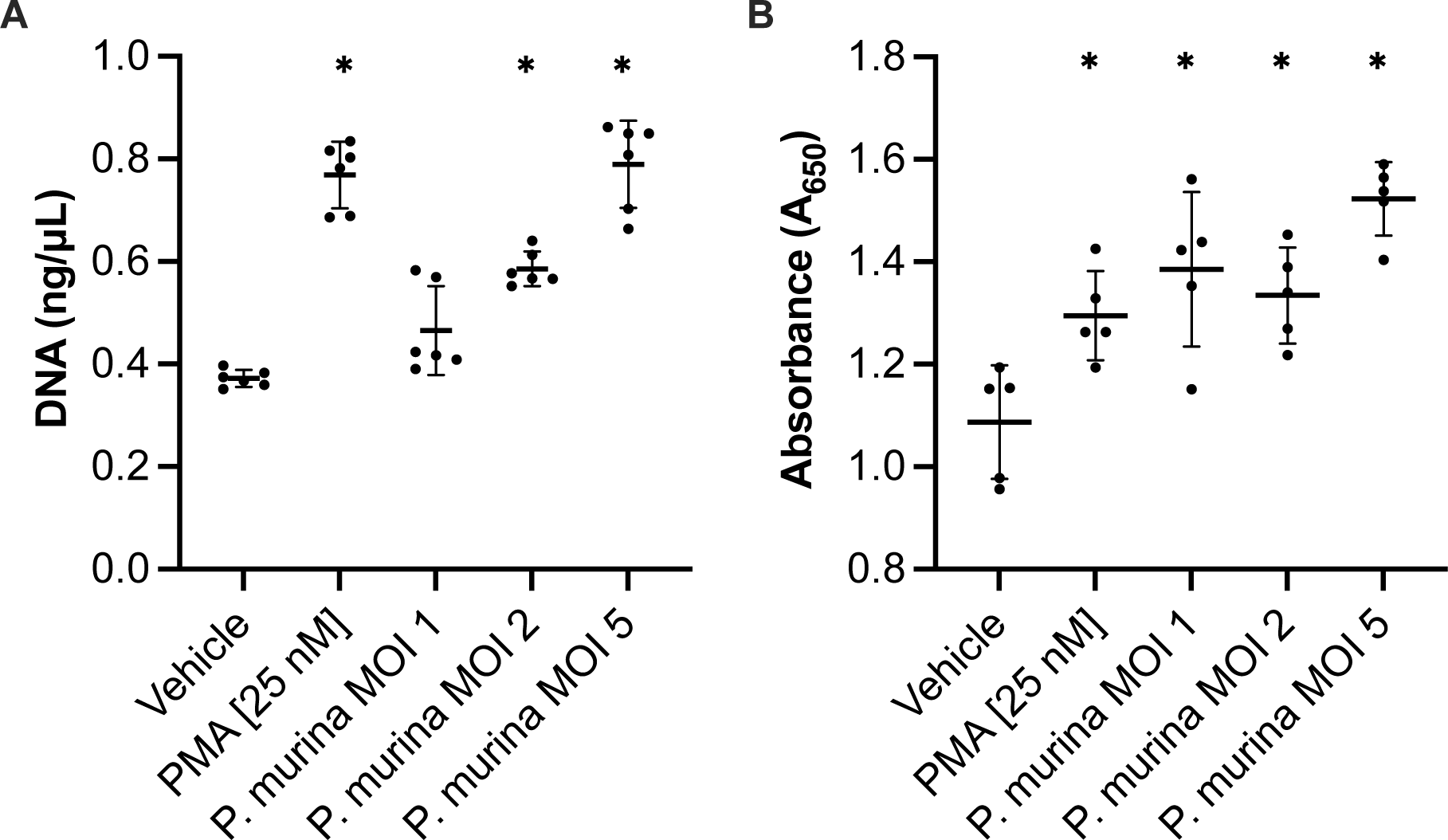
*P. murina* stimulates NET production in bone marrow-derived neutrophils. Bone-marrow derived neutrophils were stimulated with vehicle, PMA (25 nM; positive control), and *P. murina*. (A) Culture supernatant was assessed for extracellular DNA released from neutrophils. (B) Culture supernatant assessed for MPO-DNA complexes by ELISA. One-way ANOVA. *, p<0.05 compared to vehicle.

Immunofluorescence analysis showed the presence of NET structures, as seen by expelled DNA decorated with NE, CitH3, and MPO (Figure 5). These data indicate that *P. murina* can directly stimulate the production of NETs *in vitro*.

**Figure 5.**
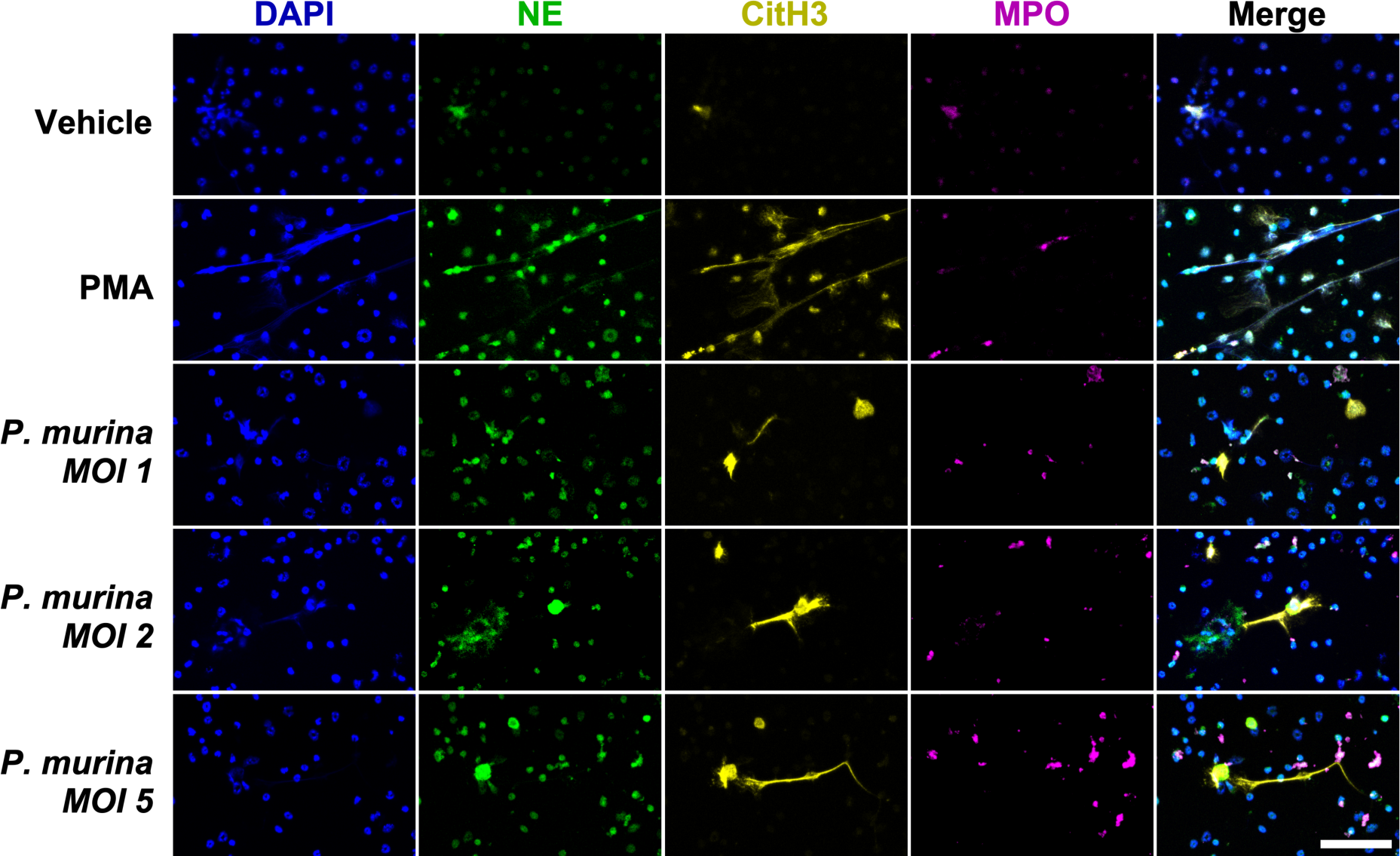
*P. murina* directly stimulates NET production in bone marrow-derived neutrophils. Immunofluorescence of the bone marrow-derived neutrophils stimulated with vehicle, PMA [25 nM], or *P. murina*. The NET production was seen in PMA-and *P. murina*-treated neutrophils as shown by neutrophil elastase (green), citrullinated histone H3 (yellow), MPO (magenta) expression. DAPI, blue. Scale, 50 µm.

### NETs are not detrimental to *P. murina* viability

NETs are known to play a role in pathogen trapping and killing. Given that *P. murina* can directly stimulate the production of NETs in neutrophils *in vitro*, we investigated whether NETs were toxic to *P. murina*. After treating *P. murina* with NETs isolated from PMA-treated neutrophils, we observed no loss of viability in *P. murina* (Figure 6). These data indicate that NETs did not negatively impact the viability of *P. murina*.

**Figure 6.**
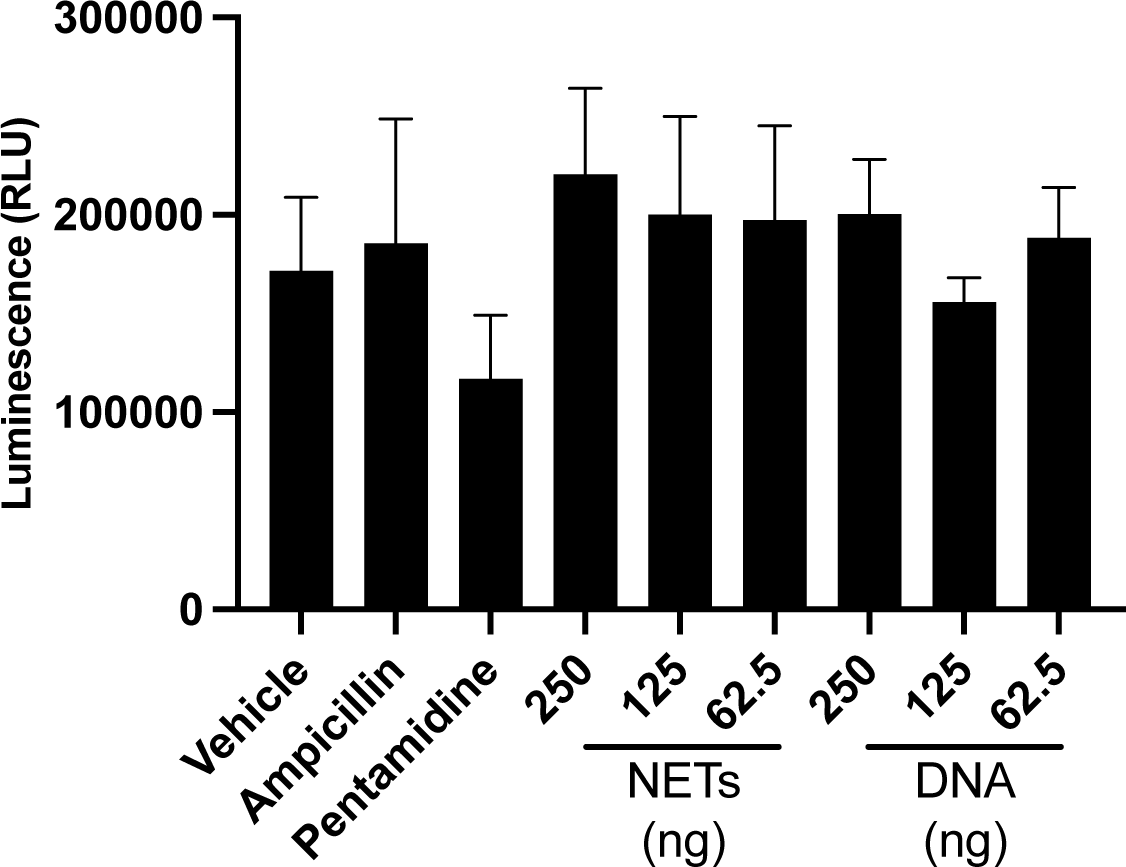
NETs are not detrimental to *Pneumocystis* viability. *P. murina* (n = 3; 5 x 10^7^ nuclei) were inoculated into 96-well plates (Costar 3548, Corning, New York) . Isolated NETs from PMA (25 nM)-stimulated neutrophils or Salmon sperm DNA samples, used as control for non-complexed DNA, were added to the wells. Plates were incubated at 5% CO_2_, 37°C. After 24 hours, 100 μL samples were transferred to opaque white plates (USA Scientific, Ocala, FL) and assessed for ATP content using ATPlite (Perkin-Elmer, Waltham, MA). No significance was seen in NET- or DNA-treated samples.

## DISCUSSION

*P. jirovecii* pneumonia remains a significant concern, particularly in patients with HIV/AIDS and those undergoing immunosuppressive treatments for conditions like hematological cancers. Immunosuppressed hosts with PjP experience an influx of various immune cells, including T lymphocytes, polymorphonuclear neutrophils, and other leukocytes, resulting in profound inflammation and considerable morbidity and mortality. Traditionally, the focus of research on host responses to *Pneumocystis* has been centered around macrophage responses and role in pathogen clearance. However, our findings highlight the overlooked roles of neutrophils and the NLRP3 inflammasome in the host immune response during *P. murina* infection.

Neutrophils, despite their abundance in infected lungs, have been largely overlooked in the context of immune response to *Pneumocystis*. These cells play a vital role in the inflammatory response during *P. murina* infection, as indicated by the increased expression of NLRP3 inflammasome-related genes observed in the lungs (Fig. 2). Moreover, our study identified NE and Citrullinated Histone H3, NET markers of NETosis, in the lungs of *P. murina*-infected mice (Fig. 3), indicating that this process was active during *P. murina* pneumonia. Our *in vitro* experiments further demonstrated that *P. murina* can directly stimulate NETosis in isolated neutrophils (Fig. 4, 5).

Neutrophils are recognized as a critical line of defense against various pathogens. Although previous studies had questioned their significance in *Pneumocystis* infections and the clearance of *Pneumocystis*, our results indicate that neutrophils may contribute to the host’s inflammatory response by initiating NLRP3-inflammasome assembly and undergoing NETosis. The increased expression of NLRP3 inflammasome-related genes and the presence of NETs suggest that neutrophils are actively involved in the immune response against *P. murina*.

Importantly, uncontrolled inflammasome activation and NETs can drive excessive inflammation and tissue damage. Previous research has demonstrated that NETs can exacerbate lung injury and disrupt barrier function in mice, while disruption of NET formation or NET degradation led to decreased tissue damage and increased survival in mice (36–38). Our study highlights the importance of understanding the role of neutrophils and NET formation during *Pneumocystis* infection, as they may be driving increased inflammation, immune cell recruitment, and tissue damage.

Here we showed that NETs did not impact *P. murina* viability *in vitro* (Fig. 6). This aligns with the previous study by Swain et al. that demonstrated the lack of fungal clearance by neutrophils (25). Alternatively, *Pneumocystis* may be utilizing extracellular DNA as a scaffold to form a biofilm (39) similar to fungal pathogens like *Candida albicans* and *Aspergillus fumigatus*, where DNA serves as a crucial component in biofilm matrices (40–42). Additionally, the integrity of these biofilms can be disrupted by DNase, potentially increasing susceptibility to antifungal therapies (42, 43)

In conclusion, our research provides valuable insights into the role of neutrophils, particularly their involvement in NLRP3 inflammasome activation and NETosis, in the host immune response during *P. murina* pneumonia. By elucidating the participation of neutrophils and their ability to form NETs in response to *P. murina*, we expand our understanding of the intricate inflammatory dynamics at play during *Pneumocystis* infection. These findings offer potential avenues for therapeutic interventions aimed at modulating the immune response in individuals at risk of or affected by PjP. Further investigations are warranted to delve into the precise mechanisms underlying neutrophil-mediated inflammation, inflammasome activation, and NETosis in PjP, as well as to explore the potential therapeutic strategies that may arise from these discoveries.

## MATERIALS AND METHODS

### Animals

Male C3H/HeNCrl (5 weeks old; Charles River, Raleigh, NC) mice were housed under barrier conditions with autoclaved food and bedding in sterilized cages equipped with sterile microfilter lids. Mice were immunosuppressed with dexamethasone (4 mg/L) in acidified drinking water, available *ad libitum*. The mice were infected by co-housing with *P. murina*-infected mice. Infection was allowed to progress for 5 weeks (to represent a moderate infection; MI) and 7 weeks (high infection; HI). Time-matched immunosuppressed, uninfected mice were used as controls (moderate and high control; MC and HC). Mice were euthanized humanely and their lungs removed for quantification of fungal burdens (n=3) or the lungs were flash frozen in liquid nitrogen, ground using a mortar and pestle, then stored at -80°C for RNA sequencing (n=5). To quantify fungal burden, lungs were homogenized in PBS using gentleMACS (Miltenyi Biotec, Auburn, CA, USA), then stained with a modified Diff-Quik staining (44) to visualize the nuclei for microscopic enumeration. The microscopic counts were log transformed, and values were compared by the one-way analysis of variance (ANOVA).

These studies were performed in accordance with the Guide for the Care and Use of Laboratory Animals, 8th ed. (National Academies Press, Washington, DC, USA), in AAALAC-accredited laboratories under the supervision of veterinarians. In addition, all procedures were conducted in compliance with the Institutional Animal Care and Use Committee at the Veterans Affairs Medical Center, Cincinnati, OH, USA.

### RNA sequencing

The following steps were performed at UC Genomics, Epigenomics, and Sequencing Core, Department of Environmental Health, University of Cincinnati, Cincinnati, OH. RNA was extracted from lung tissue using mirVana™ miRNA Isolation Kit (Invitrogen, Carlsbad, CA). The Ribo-Zero Gold (Human/Mouse/Rat) and (Yeast) kit (Illumina, San Diego, CA) were used to deplete rRNA using 300 ng total RNA as input. The isolated RNA was RNase III fragmented and adaptor-ligated using PrepX mRNA Library kit (Takara, Mountain View CA), then converted into cDNA using Superscript III reverse transcriptase (Lifetech, Grand Island, NY). Resulting cDNA was purification using Agencourt AMPure XP beads (Beckman Coulter, Indianapolis IN). Barcode index was added using a universal and index-specific primer with PCR to each ligated cDNA sample and the amplified library was enriched by AMPure XP beads purification.

The quality and yield of the purified library was analyzed by Bioanalyzer (Agilent, Santa Clara, CA) using DNA high sensitivity chip. Libraries were quantified by qPCR measured by Kapa Library Quantification kit (Kapa Biosystems, Woburn, MA) using ABI’s 9700HT real-time PCR system (Lifetech, Grand Island, NY). Libraries at the final concentration of 12.0 pM were clustered onto a flow cell using Illumina’s TruSeq SR Cluster kit v3 and sequenced for 50 cycles using TruSeq SBS kit on Illumina HiSeq system.

### RNAseq analysis

Raw reads were trimmed for quality and barcode removal using fastp (45). Trimmed reads were aligned to the *P. murina* genome (accession GCF_000349005.2) and quantified using Salmon (46). DEseq2, using an absolute fold change >1.5 and FDR adjusted p-value of 0.05, was used to identify differentially expressed genes between groups (47). Ensemble of Gene Set Enrichment Analyses (EGSEA) was performed to identify pathways with enriched gene expression with an absolute fold change >1.0 and FDR adjusted p-value of 0.05 (48).

### Immunofluorescence

Lung tissue (n = 3) was fixed in neutral buffered formalin for 24 hours at room temperature. The tissue was then dehydrated and stored in 70% ethanol. Fixed lungs were embedded with paraffin then sliced at 10 um and mounted on positive charged slides.

Neutrophils (4 x 10^4^; performed in triplicate) were grown on 18 mm glass coverslips (Electron Microscopy Sciences; Hatfield, PA, USA) in a 12-well dish (CytoOne; Ocala, FL, USA). After NETosis experiments, cells were fixed in 3.7% formaldehyde in PBS for 15 minutes and permeabilized with 0.1% Triton-X for 10 minutes.

Lung or neutrophil samples were blocked in 10% goat serum for 1 hour, then incubated with anti-Neutrophil Elastase-AlexaFluor 488 (1:100; Santa Cruz Biotechnology, Dallas, TX), anti-Citrullinated Histone H3 (1:100; Abbomax, San Jose, California), or anti-Myeloperoxidase (1:100; Santa Cruz Biotechnology, Dallas, TX) in blocking buffer for 1 hour. Secondary goat anti-rabbit antibody conjugated with AlexaFluor 594 (1:1000; Invitrogen, Carlsbad, CA) and goat anti-mouse IgG antibody conjugated with AlexaFluor 647 (1:1000; Invitrogen, Carlsbad, CA) in blocking buffer were added and incubated for 2 hours in the dark. Cells were washed between incubations with 0.1% Tween 20 in PBS (PBST) three times for 15 minutes. DNA was stained with DAPI (1 ug/mL) Images taken on a Nikon A1 Confocal Microscope at Cincinnati Children’s Hospital Medical Center or on a Leica Stellaris confocal microscope at the University of Cincinnati Live Microscopy Core.

### Neutrophil isolation

Bone marrow was obtained from the humerus, femur, and tibia of immunocompetent C3H/HeNCrl male mice (n= 3 groups of pooled bone marrow from 2 mice). Red blood cells were lysed in 0.08% ammonium chloride for 10 minutes on wet ice, then washed with RPMI 1640 (Gibco; Grand Island, NY, USA) twice. Neutrophils were isolated by positive selection using Anti-Ly-6G MicroBeads UltraPure (Miltenyi Biotec; Auburn, CA, USA) and resuspended in RPMI 1640.

### NETosis assay

Neutrophils (1 x 10^6^) were plated onto a 24-well dish (CytoOne; Ocala, FL, USA). Cells were then treated with either RPMI 1640 vehicle, phorbol myristate acetate (PMA; 25 nM in RPMI 1640; Millipore; Burlington, MA, USA), or *P. murina* at multiplicities of infection (MOI) of 1, 2, or 5. Cells were incubated at 37°C 5% CO_2_ for 4 hours to allow NETosis to occur (49). Neutrophils were then gently washed twice with PBS. Cells were resuspended in 500 µL PBS by using a cell scraper to lift the neutrophils from the plate surface, then centrifuged at 300 x g to separate cell debris from NET material. These experiments were performed in triplicate using isolated neutrophils from 3 independent groups, as described above. The NET-containing supernatant were collected for downstream analysis.

### Quantification of extracellular DNA

Extracellular DNA release was quantified from the supernatant of NETosis-stimulated neutrophils using QuantiFluor dsDNA system (Promega; Madison, WI) per manufacturer instructions. Briefly, samples and dsDNA standards were stained with QuantiFlour dsDye then fluorescence (504nm_Ex_/531nm_Em_) was measured using a BioTek Synergy HTX plate reader.

### Detection of MPO-DNA complexes

NET release from the supernatant of NETosis-stimulated neutrophils was determined by detecting complexes of DNA and MPO. High binding 96-well plate (Corning 2592; Kennebunk, ME, USA) was coated overnight with capture antibody, anti-MPO (1:500; Invitrogen; Carlsbad, CA, USA), in 50 mM carbonate buffer, pH 9.4 at 4°C. After that time, plates were washed three times with PBS with 0.1% Tween 20 (PBST). Wells were blocked with StartingBlock PBS Blocking Buffer (Thermo Scientific; Rockford, IL, USA) for 2 hours at room temperature, then washed 3 times with PBST. Cell supernatant from each treatment was diluted 1:10 and 100 μL was added to the wells and incubated at 4°C for 24 hours, then washed 3 times in PBST. Detection antibody, Anti-DNA-HRP (1:100; Zymo Research; Irvine, CA), was added to the plate and incubated for 24 hours. After washing 3 times in PBST, colorimetric signal was detected using TMB (Thermo Scientific; Rockford, IL, USA). Absorbance (650 nm) was measured using a BioTek Synergy HTX plate reader.

### NET toxicity

Isolated NETs from the supernatant of PMA-stimulated neutrophils were collected and quantified as shown above. Salmon sperm DNA was used as a control for non-complexed DNA. *P. carinii* (5 x 10^7^ nuclei) were inoculated into 96-well plates (Costar 3548; Corning, NY, USA). PBS vehicle, DNA, or NET samples were added to the wells in triplicates. Plates were incubated at 5% CO_2_, 37°C. After 24 hours, 100 μL samples were transferred to opaque white plates (USA Scientific; Ocala, FL, USA) and assessed for ATP content using ATPlite (Perkin-Elmer; Waltham, MA, USA).

## DATA AVAILABILITY STATEMENT

Fastq files have been deposited to NCBI Sequence Read Archive under BioProject Accession: PRJNA1076530.

## ACKNOWLEDGMENTS

This work was supported by HHS NIH R01HL146266 and VA I01 BX004441. MTC is a Senior Research Career Scientist supported by IK6BX005232 Department of Veterans Affairs. We thank Margaret Collins for her assistance with the ELISA. Confocal microscopy performed at the University of Cincinnati Live Microscopy Core was supported by NIH S10OD030402.

